# Microscopic Origins of Flow Activation Energy in Biomolecular Condensates

**DOI:** 10.1101/2024.09.24.614801

**Authors:** Sean Yang, Davit A Potoyan

## Abstract

Material properties of biomolecular condensates dictate their form and function, influencing the diffusion of regulatory molecules and the dynamics of biochemical reactions. The increasing quality and quantity of microrheology experiments on biomolecular condensates necessitate a deeper understanding of the molecular grammar that encodes their material properties. Recent reports have identified a characteristic timescale related to network relaxation dynamics in condensates, which governs their temperature-dependent viscoelastic properties. This timescale is intimately connected to an activated process involving the dissociation of sticker regions, with the energetic barrier referred to as flow activation energy. The microscopic origin of activation energy is a complex function of sequence patterns, component stoichiometry, and external conditions. This study elucidates the microscopic origins of flow activation energy in single and multicomponent condensates composed of model peptide sequences with varying sticker and spacer motifs, with RNA as a secondary component. We dissected the effects of condensate density, RNA stoichiometry, and peptide sequence patterning using extensive sequence-resolved coarse-grained simulations. We found that flow activation energy is closely linked to the lifetime of sticker-sticker pairs under certain conditions, though the presence of multiple competing stickers further complicates this relationship. The insights gained in this study should help establish predictive multiscale models for the material properties and serve as a valuable guide for the programmable design of condensates.

## I. INTRODUCTION

Organizing biomolecules and biochemical reactions within the crowded environments of bacterial and eukaryotic cells frequently involves forming membraneless condensate phases. [1–4] The material properties of these condensates provide a critical link between the biomolecular composition of condensates and their cellular functions [5–8]. There is a growing appreciation that the viscosity is a functionally critical feature of condensate phases, the misregulation of which can lead to the transition of fluid condensates to solid-like pathological states [9–11].

While the rheology of biomolecular condensates is a relatively new field of study, the rheology of polymers and soft materials has a long and rich history. The viscosity of polymers has been extensively studied, with many theories explaining how concentration, chain length, and monomer type contribute to polymers’ viscosities and flow dynamics [12– 14]. One of the universal features of the viscosity of polymers is their exponentiation dependence on temperature following the glass transition temperature. There exists a large body of literature since the 1930s showing that the temperature dependence of viscosity for a wide variety of simple and complex fluids obeys an Arrhenius-type equation [15–17]. However, the microscopic underpinning of activation energy is not well understood even for simple polymers and is often regarded as a phenomenological fitting parameter [18, 19]. In a recent report, we have shown that the temperature dependence of viscosity in the model peptide nucleic acid condensates for room temperatures is well-described Arrhenius-like behavior with pronounced sequence dependence [20]. Because of the initiated relationship between viscosity and dynamics of flowing and fusing of condensate phases, we have called the pre-exponential factor in temperature-dependent viscosity the flow activation energy. Following a transition rate theory interpretation of viscosity in condensed phases, the flow activation energy represents the energy barrier for reconfiguring the condensate fluid network [21–23]. Our previous experiments have led us to hypothesize that flow activation energy may be directly linked to the rate-limiting dissociation of sticker motifs present in most phase-separating proteins and nucleic acids [20]. Further support for this idea comes from detailed atomistic simulations between sticker motifs, which show a strict correlation between dissociation-free energy and condensate viscosity.

The present study aims to explore the microscopic origin of activation energy in model homotypic and heterotypic condensates with dominant and competing sticker-sticker interactions. By employing coarse-grained models of proteins and nucleic acids, we have systematically investigated the relationship of activation energy on molecule-level properties, e.g., sequence variants, compositions, and local density caused by global external factors. For the simple-component condensates, we have chosen the GY23 peptide and several of its variants, and for the two-component condensates, we have chosen positively charged pentapeptide repeat motifs and negatively charged single-stranded RNA. Both of these systems have corresponding experimental studies to which we compare our computational results and make new predictions [20, 24, 25].

We find a relationship between activation energy and characteristic relaxation times of condensate phases. By varying the hydrophobicity of the peptide sequence, we find a strong correlation with activation energy in homotypic condensates. On the other hand, we find that mutations that localize charged residues don’t significantly impact the activation energy for two-component condensates, which is different from the previous results on single-component condensates. The variance of pressure and stoichiometry change the value of viscosity and contact lifetime but affect activation energy little. This shows a different micro-mechanism on local density and reconfiguration energy barrier.

The study elucidates the microscopic origins of viscosity in biomolecular condensates, linking the activated processes of sticker dissociation to a mesoscopic quantity measurable through microrheology experiments. This established framework, along with additional data on the flow activation energies of condensates, is expected to enhance our fundamental understanding of the sequence evolution of phase-separating peptides. Moreover, it will assist in the design of novel condensate phases with tunable material properties.

## II. METHODS

### A. Description of models

We have employed a one-bead hydrophobicity scale (HPS) [26] and multi-bead Hybrid-resolution (HyRes) coarsegrained (CG) models [27] to calculate the viscosity and activation energies of GY23 peptide sequences. After validating the calculations by reproducing the sequence dependence of activation energies observed experimentally, we examined two-component condensates using a one-bead model. The two-component system comprises single-strand RNA and peptides with 40 and 25 beads, respectively. The beads in the RNA chain represent uracil-base nucleotides (U), and the beads in the peptide represent amino acids with a limited alphabet consisting of typical sticker and spacer residues R, G, and P. The non-bonded interactions between residues *i, j* are modeled via the Ashbaugh-Hatch potential, and salt-screened electrostatic interactions are modeled via the Debye-Huckel potential:

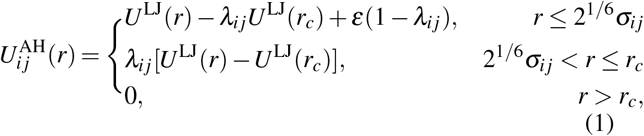

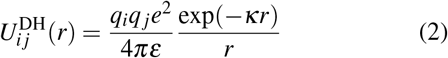

where *ε* = 0.8368 kJ/mol, *r*_*c*_ = 4 nm, and *U* ^LJ^ is the Lennard-Jones potential *U* ^LJ^(*r*) = 4*ε*[(*σ*_*ij*_*/r*)^12^ *-* (*σ*_*ij*_*/r*)^6^]. *σ*_*ij*_ = (*σ*_*i*_ + *σ*_*j*_)*/*2 and *λ*_*ij*_ = (*λ*_*i*_ + *λ*_*j*_)*/*2 are arithmetic averages of monomer size and hydrophobicity value, respectively. The pair potential with *λ* = 0 consists of only the repulsive term equivalent to the Weeks-Chandler-Andersen functional form [28]. We use *σ* values of amino acids from van der Waals volumes [29]. *σ* of nucleotide U is set as 0.817 nm. We use *λ* values of amino acids from the recently proposed CALVADOS 2 parameters [30]. *λ* of nucleotide U is set as -0.027 [31]. In the Debye-Huckel potential, *q* is the charge number of particles, and 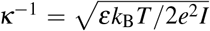 is the Debye length. In our simulations, the ionic strength of the electrolyte *I* = 0.1 nm^*-*3^. The temperature-dependent dielectric constant used in our simulations has the following empirical relation: *ε* (*T*)*/ε*_0_ = 5321*/T* + 233.76 *-*0.9297*T* + 1.41710 × 10^*-*3^*T* ^2^ *-* 8.29210 × 10^*-*7^*T* ^3^ All coarse-grained beads are connected via bonded interactions modeled by harmonic potentials 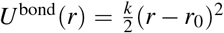 where *k* = 1000 kJ/mol/nm^2^, *r*_0_ = 0.5 nm for RNA [31] and *r*_0_ = 0.38 nm for peptide. We have not employed any angular or dihedral potential in the simulations.

In the HyRes model, we set the temperature and ionic strength (*I*) to match those used in the HPS model, with other parameters adopted from [32].

### B. Molecular simulations

We have used OpenMM (v7.7) for single-component condensate simulations. The number of chains in the simulation box was set at *N*_pep_ = 800. The initial length of the cubic box was set at 30 nm, which is much larger than the equilibrium size. The box was then shrunk in the subsequent *NPT* simulations. After energy minimization, we used the *NPT* ensemble with a timestep 0.01 ps, to equilibrate all systems using additional 2 ×10^7^ steps. Finally, we fix the box size as the average in *NPT* simulations and carry out additional *NVT* simulations for 5 ×10^7^ steps.

By the Green-Kubo (GK) relation, the shear stress relaxation modulus *G*(*t*) can be determined by computing the auto-correlation of any of the off-diagonal components of the stress tensor. A more accurate expression can be obtained as our isotropic system [33, 34]

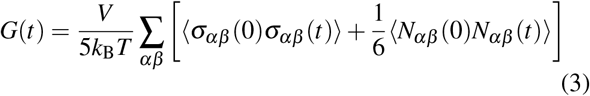

where *N*_*αβ*_ = *σ*_*αα*_ *σ*_*ββ*_ is the normal stress difference and the summation contains (*αβ* = *xy, yz, xz*). Then, the shear viscosity can be straightforwardly calculated by integrating the shear stress relaxation modulus in time. To avoid the typical noisy nature of *G*(*t*) at long time scale, we fitted *G*(*t*) to a series of Maxwell modes *G*_M_(*t*)= ∑_*m*_*G*_*m*_exp (−*t/τ*_*m*_) which are equidistant in logarithmic time [34, 35]. Therefore, viscosity is obtained by

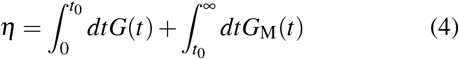

The division time *t*_0_ is chosen so that all intra-molecular oscillations of *G*(*t*) have decayed, and the function becomes positive after *t*_0_. Starting from the equilibrium, we run *NVT* simulations for 10^5^ steps and record trajectories of all beads every 50 steps. We recorded the stress tensor matrix every 10 steps for evaluating short-time integral and 100 steps for evaluating long-time integral (see Eq. (4)) in the *NVT* simulations. A similar equilibration protocol is used for two-component systems, which include a variable number of peptides, *N*_pep_ and RNA, *N*_RNA_ chains. The total number of the CG monomers was kept at about 2 × 10^4^ to eliminate the finite-size effect.

We have also used non-equilibrium oscillatory shear simulations for two-component systems to ascertain the accuracy of viscosities in heterogeneous mixed condensate states.

For non-equilibrium simulations, we have used the LAMMPS package to impose a constant rate of shearing strain 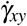 to the whole system in *NVT* ensemble for over 10^7^ steps. Only the later 50% data is used for calculating viscosity. The stress tensor *σ*_*xy*_ is recorded every 10 steps. The viscosity *η* from oscillatory shear simulations is obtained via the following equation by taking the limit when 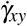 goes to zero

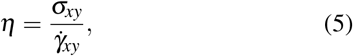

We set 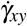 as 2 ×10^*-*5^ to 2 ×10^*-*4^ ps^*-*1^ which are slow enough to get viscosity independent of 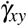. We reduce error by reducing the average viscosity with different 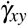.

## III. RESULTS AND DISCUSSION

### A. Single-component condensates

The GY23 peptides are convenient model systems for investigating sequence-dependent material property trends because of their great sensitivity to point mutations [24, 32]. Recently, viscosities of GY23 peptides have been measured by dynamic light scattering experiments [36]. Additionally, simulations of GY23 sequences employing the recently developed HyRes model [32] have been shown to capture the thermodynamics of phase equilibria and *c*_*sat*_ values. Following the sequences used in the experiments, we first set out to capture the viscosity dependence of single-component GY23 condensates in its wild-type (WT) form and several mutations. The mutations are grouped into decreasing hydrophobicity (F9A, F9L, and H12K) and increasing hydrophobicity (A7F and A7A10F). Their sequences and hydrophobicity difference (∆ *λ*) are shown in the table I. The temperatures of simulations for each mutation range from 75% to 95% of the critical temperature determined by coexistence simulations [26, 31, 34] (critical temperatures are detailed in SI-B). We find the monomer number density decrease with increasing temperature (see Fig.1(a)).

**TABLE I.**
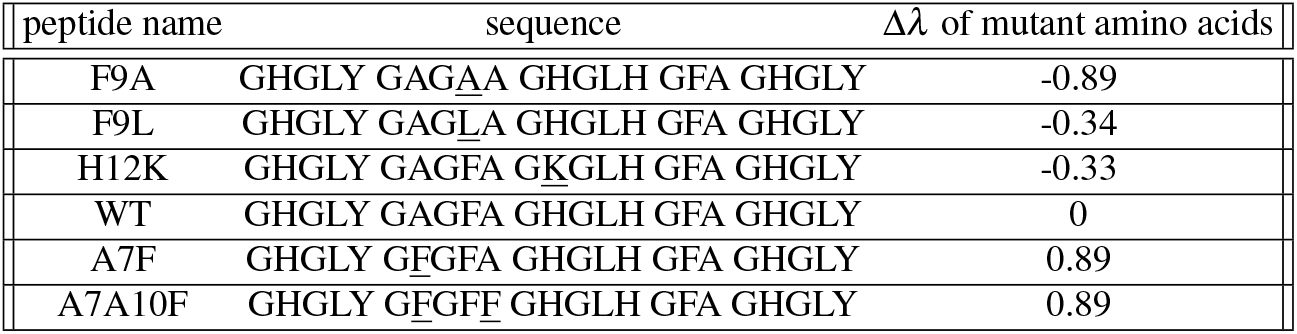
Sequences of GY23 variants. Amino-acid mutations are underlined in sequence and their hydrophobicity changes ∆ *λ* are shown.

For each variation of GY23, we observed that increasing temperature leads to an exponential decrease in the viscosity, which can be fitted with an Arrhenius-like exponential function, as shown in Fig. 1(b)

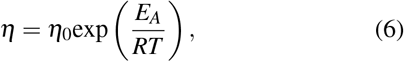

where *R* is the universal gas constant, *η*_0_ is the viscosity prefactor and *E*_*A*_ is called flow activation energy [20]

**FIG. 1.**
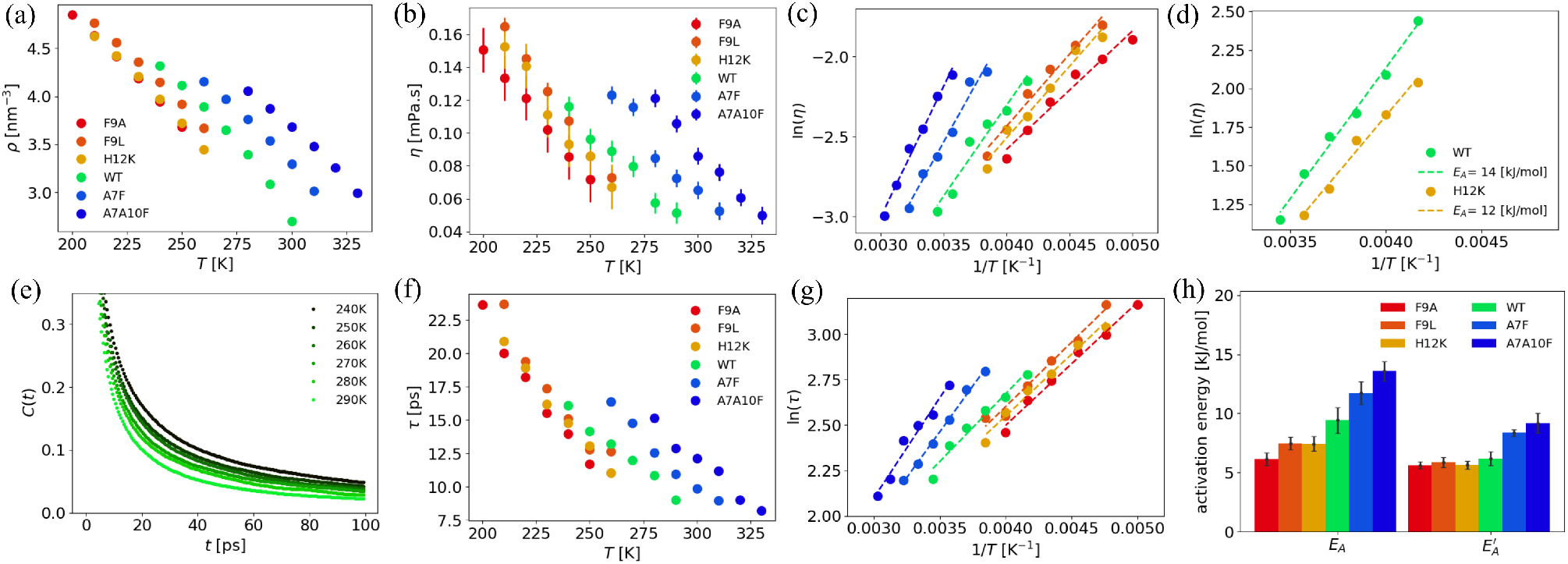
Viscosity and contact time analysis of GY23 and its variants (a) The number density of CG monomers *ρ* as a function of *T* for different variants. (b) Viscosity *η* as a function of *T*. (c) Linear fittings of ln*η* v.s. 1*/T*. (d) ln*η* v.s. 1*/T* and activation energies from HyRes model. (e) Time correlation *C*(*t*) of WT GY23 for different temperatures. (f) Contact time *τ* as a function of *T*. (g) Linear fittings of ln*τ* v.s. 1*/T*. (h) The activation energies of viscosity and contact time from (c) and (g).

A commonly employed flow model for simple and complex fluids is based on the transition state theory, which implies the existence of energetic barriers for rearranging molecular contacts [37, 38]. The energetic barrier is quantified by flow activation energy *E*_*A*_, the microscopic origin of which, just like in chemical reactions [21], is a convoluted function of molecular structure, environment, and intra-molecular interactions. The prefactor *η*_0_ depends on local molecular packing density and is related to the frequency of barrier-crossing attempts in rate theories [39]. Despite the naive view of activated rate theory on complex dynamic processes such as a flow of complex biopolymeric materials, the phenomenological relation did prove rather helpful for gleaning insights on microscopic origins of condensate viscosities [20].

Our simulations show that tuning the hydrophobicity value *λ* of a single residue in the GY23 peptides significantly impacts condensates’ activation energy *E*_*A*_. Specficially we find that higher *λ* corresponds to larger *E*_*A*_ (see Fig.1(h)) which matches recent experimental results[20]. A similar trend is obtained by using a structurally more accurate model of peptides, the recently developed HyRes model, which could better describe the backbone and transient secondary structures of protein [27, 32]. The results also exhibit Arrhenius-like behavior and qualitatively follow the same trend as the variant from WT to H12K, as shown in Fig. 1 (d).

To gain microscopic insight into the nature of activation energy, we examined the typical relaxation timescales in fluids. Relaxation time is the characteristic time required for biomolecules to reconfigure contacts with neighboring biomolecules. The probability of reconfiguration over an energy barrier is proportional to a Boltzmann factor exp (−*E*_reconf_*/RT*). As a reciprocal of the reconfiguration rate, the relaxation time would have an Arrhenius-like form proportional to exp(*E*_reconf_*/RT*) [37, 38]. In complex bio-molecular mixtures, the relaxation time shares a mechanism similar to that of fluid, while the reconfiguration mechanism is more complicated. There is a hierarchy of relaxation time scales such as the global reputation time, local segment reconfiguration time, and dynamic binding relaxation time [40]. Following the reconfiguration mechanism, we applied the microscopic lifetime of contact *τ* defined from the time correlation of contact between particles i and j:

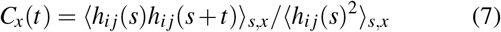

where *h*_*ij*_(*s*) is a step function to describe whether particle *i* and *j* are in contact or not. ⟨ * ⟩_*x*_ take the average in a contact group *x* of consideration. Two particles *i, j* are in contact, which means the distance *i* to *j* is less than a given value *d*. Considering the sizes of monomers are from 0.45 (P) to 0.82 (U) nm in our system, *d* = 1 nm is a proper setting to discern a dissociation of *d >* 1. The configuration around these two monomers is probably irreversible after the dissociation because other monomers will easily come to fill the gap.

The characteristic contact time is an integral of correlation function 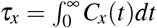. We chose *x* as all particles for single-component condensates and omitted subscript *x* = all in the following content. The time correlation functions at different temperatures have similar forms of exponential decay, as shown in Fig.1(c). We use multi-exponential function 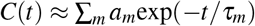 to fit these curves and calculate the integral value *τ*. For different variants of GY23, *τ* decreases as temperature increases. We found the relationship *τ* (*T*) can be fitted with an Arrhenius-like scaling function. Similar to the viscosity, we can also define a corresponding activation energy from the form

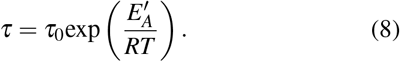

Here, 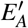 increases when mutation of higher hydrophobicity *λ* in the peptides; that tendency is the same as flow activation energy *E*_*A*_. The magnitude of 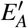 is approximately 65-90% of *E*_*A*_. This difference may result from the fixed contact distance of *d* = 1 nm used to define Eq.(7)’s contact lifetime for different densities *ρ*. This approach contrasts with certain bond-exchange simulations, which treat these factors differently. [18, 40].

### B. Two-component condensates

To better understand the microscopic origins of flow activation energy in biomolecular condensates, we next turn to a prototypical two-component condensate comprising a peptide with sticker and spacer residues and an unstructured single-stranded RNA. Recent microrheology experiments have characterized these systems, demonstrating the programmable nature of peptide-RNA condensates [20]. In these two-component systems, the RNA is fixed in length and carries a negative charge (we use 40 uridines U_40_ in the simulations), while the peptide has a net positive charge due to sticker arginine residues interspersed with spacer glycines or prolines with varying hydrophobicity. The size of CG monomers of RNA and peptide are *σ* = 0.82 and *σ* = 0.45 *-* 0.65, respectively. As a result, the radial distribution functions *g*(*r*) of monomer pairs exhibit two distinct master curves corresponding to RNA-peptide and peptide-peptide pairs. This distinction reveals the configurational differences between RNA-peptide condensates and GY23 condensates (See SI-B). The temperature was consistently maintained below the critical temperature and above the glass transition temperature to ensure uniformity throughout the system [34] (critical temperatures are provided in SI-B). We measured the viscosity of peptide-RNA condensates using the constant-shearing method (detailed in Methods B-2). Compared to the GY23 condensates, RNA-peptide condensates contain a higher fraction of charged monomers, leading to a more complex interplay between electrostatic and hydrophobic interactions. We demonstrate that RNA-peptide condensates exhibit a weaker correlation between density and viscosity *η* (*ρ*) compared to GY23 (see SI-C) This interplay also results in a distinct preference for contacts among different CG monomers, prompting us to conduct detailed statistics of RNA-peptide condensates outlined in the next section.

#### 1. Contact statistics for RNA-peptide system

The charge ratio of peptides significantly affects the physical properties of bio-condensates. For instance, the peptide (PPRPP)_5_ has a charge number of *Q* = 5, and we set *m*_*c*_ *N*_pep_*/N*_RNA_ = 8 to achieve charge neutrality. In contrast, (RPRPP)_5_ has a higher charge number of *Q* = 10, requiring *m*_*c*_ = 4 for neutrality. We calculated contact statistics for U_40_-(PPRPP)_5_ and U_40_-(RPRPP)_5_ condensates, with snapshots of these two condensates shown in Fig.2 (a-b). The contact ratios of all residue pairs from different chains are illustrated in Fig. 2 (c-d).

**FIG. 2.**
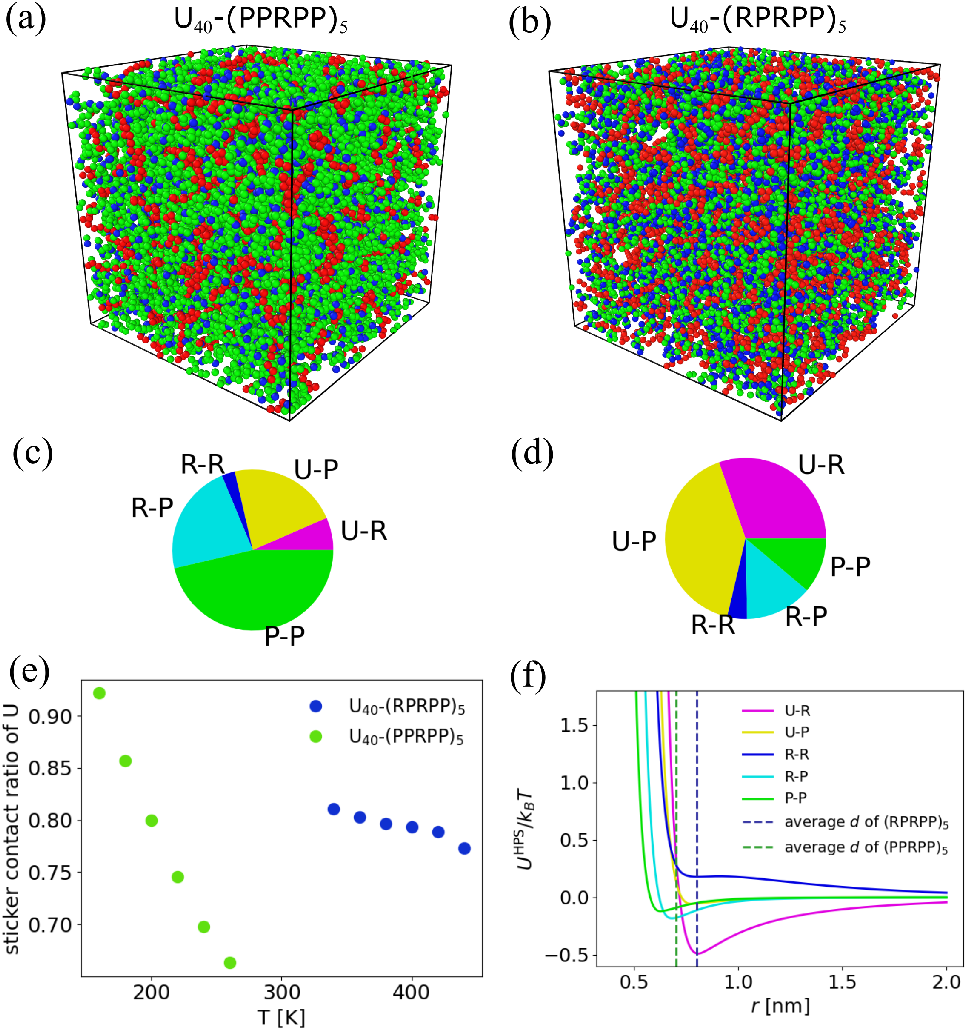
Contact statistics of RNA-peptide condensates (a-b) Snapshots of U_40_-(PPRPP)_5_ and U_40_-(RPRPP)_5_ condensates at temperatures T=240K and T=400K, respectively. The CG amino acids U,R, and P are represented by red, blue, and green spheres. (c-d) Contact ratios relative to all contact pairs for condensates (a) and (b), respectively. The U-U slice is too small to display. (e) The sticker contact ratio of residue U for different temperatures. (f) The potential of non-bonded interaction in the HPS model. Different colors represent different contacts. Two vertical lines represent two average distances of condensates in (a) and (b).

For simplicity, we classify the residue contacts into three categories based on their respective chains. Inter-chain RNA-RNA contacts (group tag ‘r-r’) correspond to residue contacts between two uracils (U-U). Inter-chain RNA-peptide contacts (group tag ‘r-p’) encompass the sum of residue contacts between uracil and arginine (U-R) and uracil and proline (U-P). Inter-chain peptide-peptide contacts (group tag ‘p-p’) include the sum of residue contacts between arginine-arginine (R-R), arginine-proline (R-P), and proline-proline (P-P). The ratio of r-r contacts to the total number of contacts is consistently very low (<0.5%) due to the strong electrostatic repulsion between RNA monomers. Therefore, r-r contacts can be ignored in subsequent calculations, allowing us to focus solely on protein-RNA and protein-protein contacts. In general, r-r contacts could be important as they contribute to hydrogen bonding and base stacking interactions; however, these interactions are not included in our CG model. As shown in Fig. 2 (c-d), p-p contacts (R-R, R-P and P-P) account for 70% of the total contacts in the low-charged U_40_-(PPRPP)_5_ condensates. In contrast, p-p contacts constitute only about 35% in the high-charged U_40_-(RPRPP)_5_ condensates.

The contact lifetime *τ*_*x*_ (*x*=rp, pp, and all, subscript ‘all’ will be omitted) varies depending on the type of contact. As shown by different symbols in Fig.4(b and f), *τ*_pp_ *> τ*_rp_ in low-charged peptide cases, whereas *τ*_pp_ *< τ*_rp_ in high-charged peptide cases. We can estimate the order of contact lifetime by the pair potential of HPS model (*U* ^HPS^ = *U* ^AH^ + *U* ^DH^) (Fig.2(f)). At the average distance (*ρ* ^*-*1*/*3^), p-p contacts are attractive, and r-p contacts are repulsive in U_40_-(PPRPP)_5_ condensates. However, r-p contacts become more attractive than p-p contacts in U_40_-(RPRPP)_5_ condensates. This corresponds to the hierarchy of contact lifetimes *τ*_pp_ *> τ*_rp_ in the former and *τ*_pp_ *< τ*_rp_ in the latter condensates. Recent studies have employed mean-field theory to estimate peptide phase separation properties and sub-region interactions based on pair potential [41, 42]. These findings may motivate a mean-field estimation of contact time and activation energy.

The concentration of contacting stickers influences the frequency of observed reconfiguration events and is directly related to the lifetime of contacts. A higher concentration of sticker contacts corresponds to slower reconfiguration time and is positively correlated with the magnitude of viscosity [43, 44]. For example, we compare the sticker contact ratio of U residues. This ratio is defined as the number of U residues within a distance of *d* = 1 nm from any R residue divided by the total number of U residues. We observe that the contact ratio decreases with increasing temperature (Fig. 2 (e)) with U_40_-(PPRPP)_5_ condensates showing a faster decline compared to U_40_-(RPRPP)_5_. This trend is similar to the behavior of *η* (*T*) and *τ* (*T*) (Fig. 4 (a-b)), as expected from the similar mechanism underlying the relaxation process.

#### 2. The effects of pressure

Biomolecular condensates forming in nucleoplasmic and cytoplasmic environments are subject to higher crowding than in solutions used in vitro experiments. The external pressure from a crowded environment tends to enhance the density of condensed phases. We conducted simulations by varying pressure from 0 to 2.5 atm for U_40_-(RGRGG)_5_ condensates to investigate how external pressure impacts activation energies. Chain number *N*_RNA_ ≈ 140 for different temperatures (details are shown in SI-A) and the number ratio *m*_*c*_ ≈ 4 to keep the whole system neutral.

For any given external pressure *p*, we find that the viscosity is still a decreasing function of temperature. The Arrhenius law also works here as well (Fig. 3 (a-b)). We note that the density increases with increasing pressure, which has a stabilizing effect on condensates (Fig. 3(c)). Unsurprisingly, increasing thermodynamic stability is accompanied by higher viscosity values, showing that pressure-increased density is a significant contributor to the viscosity of condensates. These observations match common compressible fluid characteristics because higher pressure induces higher density, which increases the energy barrier of reconfiguration in simple fluid [38]. Interestingly, however, the activation energy *E*_*A*_ extracted from the Arrhenius law remains the same (average *E*_*A*_ = 5.5 kJ/mol) regardless of the values of external pressure *p* (as shown in Fig.3(b)). That means pressure change only affects the prefactor in Eq. (6).

**FIG. 3.**
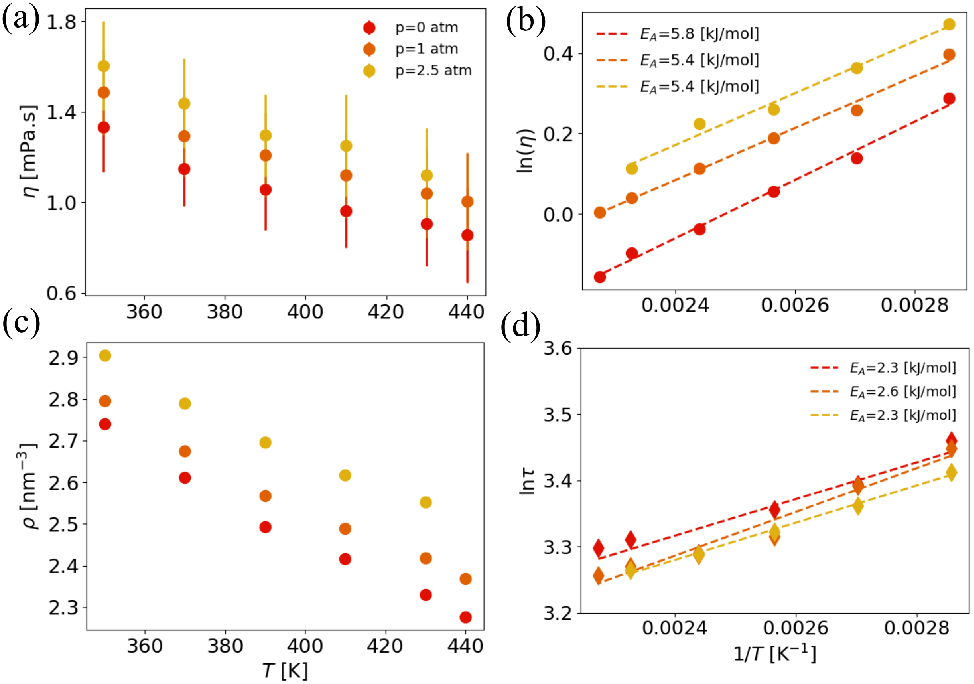
The effects of pressure on U_40_-(RGRGG)_5_ condensates. (a) Viscosity *η* as a function of *T*. (b) Linear fittings of ln*η* v.s. 1*/T*. (c) The number density of CG monomers *ρ* as a function of *T*. (d) The contact time ln*τ* as a function of 1*/T*.

**FIG. 4.**
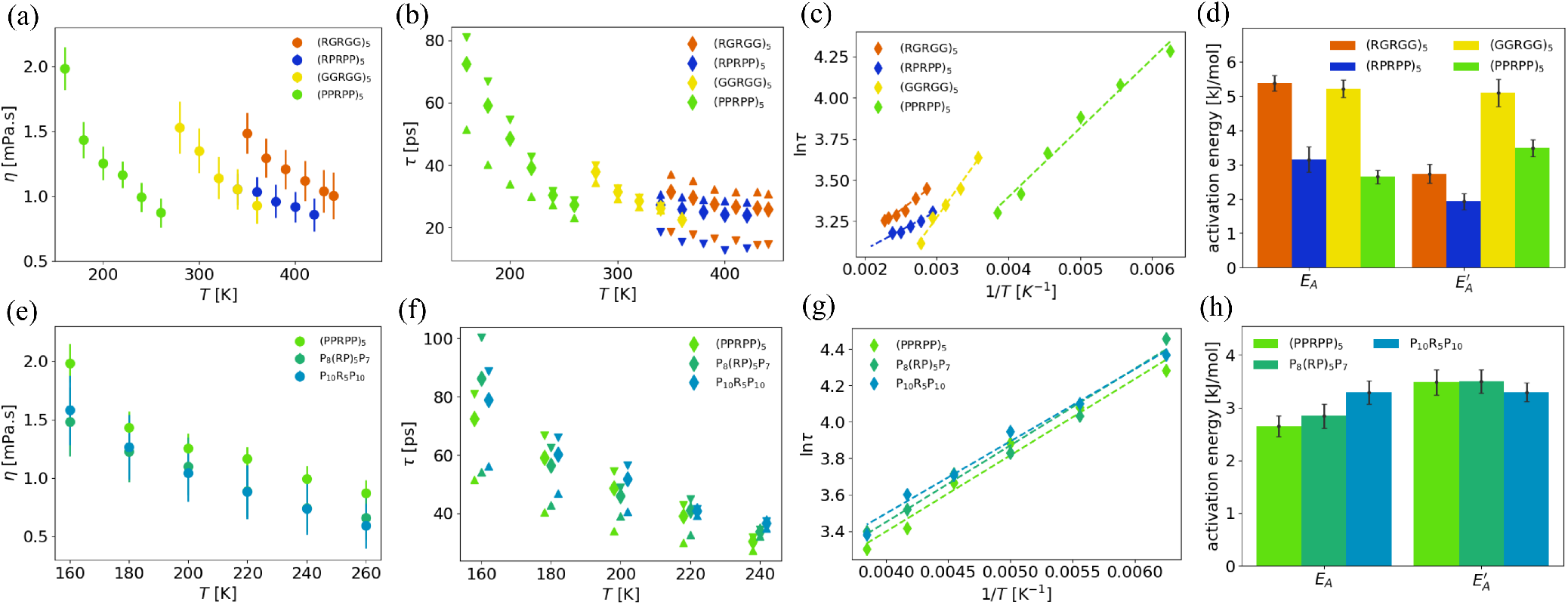
Multi-scale quantities for sequence variants. The first row (a-d) shows the results of variations of charge number *Q* and hydrophobicity *λ*. The second row (e-h) shows the results of different charge patterns. (a and e) Viscosity *η* as a function of *T*. (b and f) Contact time *τ* (*T*) calculated from different contact groups. Upward triangles, downward triangles, and diamonds represent contacts between RNA-peptide, peptide-peptide, and all averages, respectively. The temperatures in (f) are separated a bit only for display. (c and g) ln*τ* as a function of 1*/T* for average of all contacts. (d and h) Comparison of activation energy *E*_*A*_ and 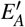.

To gain insight into the microscopic origin of pressure insensitivity of flow activation energy, we compute the characteristic contact time *τ*_*x*_ following the same method that is used for the analysis of homotypic condensates formed by GY23 peptides. The Arrhenius-like behavior of *τ* (*T*) is again observed (Fig. 3(d)), which is virtually undisturbed by external pressure with an average 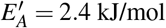, consistent with the trend of viscosity under varying pressure.

#### 3. Peptide sequence variations

To study the roles of sticker spacer patterns on the viscosity of two-component condensates, we change the sequences of peptides in three ways that systematically vary hydrophobicity, charge number, and charge localization.

First, we changed the hydrophobicity *λ* and kept the charge number *Q* the same. Replacing all G (*λ* = 0.7) to P (*λ* = 0.37), we change from (RGRGG)_5_ to (RPRPP)_5_ and (GGRGG)_5_ to (PPRPP)_5_. The simulations show both replacements decrease viscosity and corresponding activation energy *E*_*A*_. The contact time and 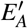 also decrease as a result of substitutions. *λ* ↓ decreases the short-range attraction in hydrophobic interaction, which affects both the prefactors and activation energies of viscosity and contact time. Our simulation results match and generalize the previous experimental observation [20].

Next, we change the charge number of a peptide chain. The amino acids R and G have similar hydrophobicity (*λ* = 0.7), but R is positively charged, and G is neutral. With 50% R replaced by G, we changed sequence (RGRGG)_5_ (*Q* = 10) to (GGRGG)_5_ (*Q* = 5). The chain number *m*_*c*_ ratio is doubled to keep the whole system neutral. We find viscosity decreases after R-G replacement with little change to *E*_*A*_, as shown in Fig. 4(a). The contact time also decreases after this replacement, but 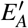 increases to more than double.

The variance of charge numbers *Q* significantly affects the component ratio of residues because it also alters the chain number ratio *m*_*c*_. Predicting the overall effect with a complicated interplay of electrostatic interaction, hydrophobicity, and density is challenging. In the observed temperature range, *Q* = 10 peptides exhibit stronger electrostatic interactions, but *Q* = 5 peptides have a higher density (see Fig. S2). This increased density counteracts the reduction in electrostatic interactions, increasing the activation energy 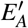.

Finally, we alter the order of charged monomers to change the charge pattern of peptides. A parameter called normalized sequence charge decorator (SCD) has been used to describe the concentration of charged monomers [45], which tends to explain some patterns of sequence-dependent material properties. We vary the sequence by sequentially grouping the charge monomer R; (PPRPP)_5_ → P_8_(RP)_5_P_7_ →P_10_R_5_P_10_. As a result of SCD variation, we find that the viscosity decreases (Fig. 4(d)), when the charge monomers are localized, with only a small change to activation energy *E*_*A*_, which indicates that activation energy is a much stronger function of charge concentration rather than patterning in our two-component system. The variation of charge pattern also doesn’t affect the contact time, leaving the 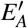 intact.

Overall, The Arrhenius-like behavior of *η* (*T*) and *τ* (*T*) always appear in the proper temperature region when we explore the variants of sequences. In agreement with prior experiments [20], the decrease on *λ* in sequence level could effectively decrease *η* and *τ* of condensates. Unlike single-component simulations from [45], the variation of charge patterning in RNA-peptide condensates does not affect *η* and *τ* a lot.

Different contact groups exhibit different contact time scales. For *Q* = 10 peptides (e.g., (RGRGG)_5_ and (RPRPP)_5_), the contact time *τ*_rp_ is greater than the average *τ* (see Fig. 4(b)). This suggests that the higher electrostatic affinity between negatively charged RNA and positively charged peptides results in a longer contact lifetime for these groups compared to others. For *Q* = 5 peptides, the contact *τ*_pp_ is larger than the average *τ*, indicating that hydrophobic attraction has a stronger influence on contact lifetime in these condensates. We can estimate this phenomenon by comparing the average amplitude of pair potentials in different condensates (see Fig.2 (f)). From the comparison of activation energies in Fig. 4(d and h), we have 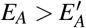 for *Q* = 10 peptides, while 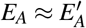 for *Q* = 5 peptides. This discrepancy may be attributed to the influence of density variations on the fixed criteria used for calculating contact lifetime in Eq.(7). If we focus on the slower contact group (*x* = rp for *Q* = 10 peptides and *x* = pp for *Q* = 5 peptides) rather than the all-average contact group, we obtain activation energies similar to 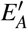. Detailed results are provided in SI-D. This indicates that the activation energy derived from our ‘average’ statistics can effectively represent the relaxation process.

#### 4. The effect of stoichiometry

In the previous section, we found that activation energy is sensitive to the concentration of charged residues and insensitive to charge patterning. Motivated by this finding, we next turn to analyze charge stochiometry dependence, which offers a glimpse into cellular regulation of the viscosity of condensates. We varied RNA concentration in two components condensates: U_40_-(GGRGG)_5_ and U_40_-(PPRPP)_5_. The RNA concentration was adjusted within a range that ensures a stable and uniform condensate phase, corresponding to *m*_*c*_ between 8 and 6. Our initial observation showed that the additional charge from RNA rapidly decreases the condensate density, *ρ* (see Fig.5(c)). In the uniform phase, the viscosity *η* and contact time *τ* decrease as *m*_*c*_ changes to 6 (see Fig.5(a-b)). Notably, both *η* (*T*) and *τ* (*T*) follow the Arrhenius law closely, with their activation energies remaining constant across the range of *m*_*c*_. This is similar to the observations from pressure variations where the prefactor *η*_0_ and *τ*_0_ is affected without much change in activation energy.

**FIG. 5.**
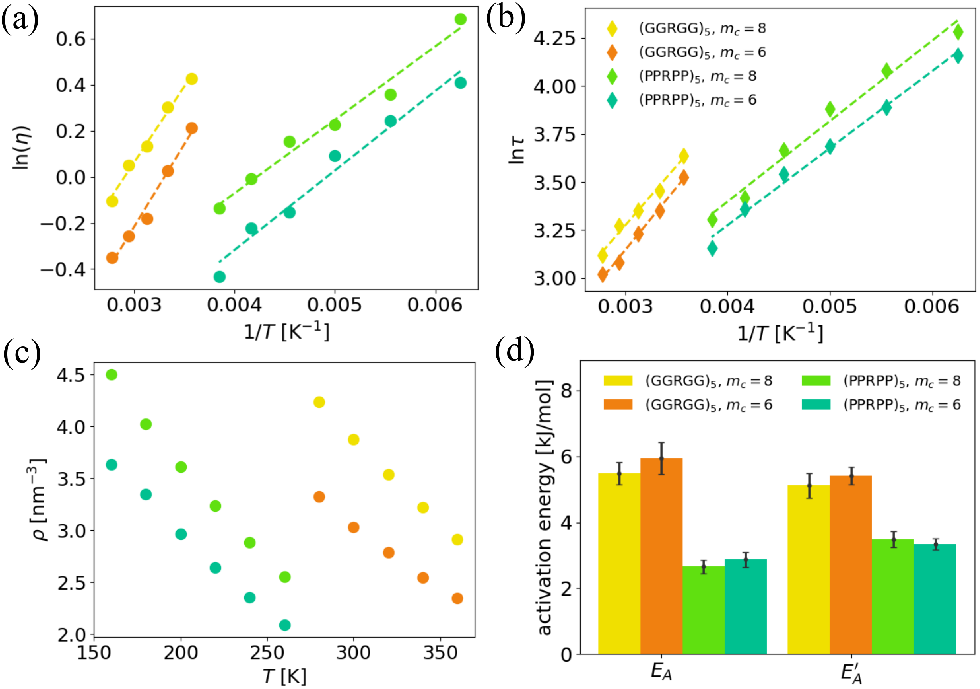
Multi-scale quantities for stoichiometry variations. (a) Viscosity as a function of *T* for different stoichiometry. (b) Contact time ln *τ* as a function of 1*/T*. (c) Number density *ρ* as a function of *T*. Comparison of activation energy *E*_*A*_ and 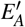.

## IV. DISCUSSION

Material properties of biomolecular condensates shape the dynamical behaviors of condensates, which directly affect their cellular functions. Recent studies have shown that the viscosity of condensates is programmable by variations in sequence, composition, and length of biomolecules. Furthermore, despite the molecular complexity, microrheology experiments have found that the temperature dependence of condensate viscosity follows the Arrhenius law. Due to the inherent molecular complexity and heterogeneity of condensate phases, it is challenging to decipher the microscopic origin of viscosity and its temperature dependence. By employing coarse-grained sequence-resolved molecular models of proteins and nucleic acids, in this work, we show that one could explain the composition and environmental effects on flow activation energy by focusing on the underlying activated processes driven by thermal fluctuations in condensates in equilibrium.

We show that the reconfiguration time scales directly impact the microscopic origin of flow activation energy in prototypical single and two-component condensates using GY23 and RNA-peptide mixtures. We observe a strong correlation between microscopic contact dynamics and viscosity by quantifying the reconfiguration time scales. Similar to the viscosity, the dissociation lifetime of sticker pairs also has Arrhenius-like behavior for both single-component and two-component condensates. Below, we outline some of the key findings of this work.

First, in both single-component GY23 peptide and two-component RNA-peptide condensates, we show that by continuously varying the overall hydrophobicity of peptide chains, one could continuously increase the activation energy *E*_*A*_ of condensates. Thus, we establish that hydrophobicity or short-range attractive forces can significantly alter energetic barriers for network reconfiguration and, hence also, flow activation energy. On the other hand, we find that the external pressure and stoichiometry only affect the prefactor *η*_0_, leaving the activation energy intact. Our findings are in harmony with reaction rate theory where prefactor *η*_0_ depends on local molecular packing density and is related to the attempt frequency for crossing energy barrier [39]. The pressure variation does change the local density, which in turn impacts the contact frequency of sticker pairs in condensates.

Unlike previous single-component simulations [45], the variance of charge segregation has little effect on viscosity and contact time in RNA-peptide condensates (Fig.4 (e)-(g)). When alternating patterns of charged stickers are kept con-stant in total number, we find that protein-RNA binding events or contacts do not change much. However, the contacts between peptides change significantly. For instance, we show that the diblock sequence produces much stronger binding between two peptides than the perfectly alternating sequence. Our two-component condensates consist of strongly charged RNA, and the interaction between RNA and peptides has more influence on the activation energy. That is the primary difference between these two condensates.

We demonstrate the existence of a hierarchy of time scales in different residue pair contacts by measuring dissociation lifetimes. We show that even minor sequence variations can affect the order of these timescales. In highly charged peptides like (RPRPP)_5_, RNA-peptide contact lifetime (*τ*_rp_) is longer than the peptide-peptide lifetime (*τ*_pp_). However, replacing (RPRPP)_5_ with (PPRPP)_5_ reverses this order: *τ*_rp_ *< τ*_pp_. Furthermore, this hierarchy of timescales can be estimated by comparing the corresponding pair potentials at the average pair distance. This may allow for future predictions of activation energy based on configuration and potentials.

In this work, we shed light on microscopic interactions that strongly correlate with flow activation energies of condensates, thereby providing a predictive link between sequence patterns of biomolecules with the material properties of resulting condensates. The insights gained in this study should help establish predictive multiscale models for the material properties and serve as a valuable guide for the programmable design of condensates.

## ACKNOWLEDGEMENTS

The authors acknowledge the support from the National Institutes of Health with grant no R35GM138243.

