## Supporting Information for "Microscopic Origins of Flow Activation Energy in Biomolecular Condensates"

### Supporting Information: "Microscopic Origins of Flow Activation Energy in Biomolecular Condensates"

##### A. Detailed settings and properties of condensates

To achieve a uniform fluid state, we set the temperatures of our condensates within a range of 70%-95% of critical temperature. We determined the critical temperatures by running slab simulations with both dilute and concentrated phases [26, 31]. The critical temperatures for GY23 and RNA-peptide condensates were obtained (as shown in table SI), and were used as reference temperatures in our subsequent simulations in the cubic box.

| peptide name | $N_{\text{pep}}$ | $N_{\text{RNA}}$ | $T_c$ [K] |
|---|---|---|---|
| F9A | 800 | / | 250 |
| F9L | 800 | / | 270 |
| H12K | 800 | / | 270 |
| H12K (HyRes) | 300 | / | 310 |
| WT | 800 | / | 300 |
| WT (HyRes) | 300 | / | 330 |
| A7F | 800 | / | 330 |
| A7A10F | 800 | / | 350 |
| $(RGRGG)_5$ | 560±80 | 140±20 | 490 |
| $(RPRPP)_5$ | 560 | 140 | 460 |
| $(GGRGG)_5$ | 640 | 80 | 370 |
| $(PPRPP)_5$ | 640 | 80 | 220 |
| $P_8(RP)_5P_7$ | 640 | 80 | 230 |
| $P_{10}R_5P_{10}$ | 640 | 80 | 240 |

TABLE SI. The chain number setting in CG simulations and critical temperatures of condensates

##### B. Radial distribution function of single and two-component condensates

For single-component condensates, we present the radial distribution function $g(r)$ for the first seven residue pairs of WT GY23, ordered by the number of pairs within a distance of 1 nm. Except for minor oscillations at short distances corresponding to monomer size, all $g(r)$ curves converge into a master curve when $r > 1$ nm. For two-componenet condensates, the size of RNA monomer $\sigma = 0.8$ nm is larger than that of peptide monomers, which range from $\sigma = 0.45$ to 0.65 nm. This results in two distinct master curves representing RNA-peptide and peptide-peptide pairs, as shown in Fig. S1(b-c). The highly-charged peptides ($U_{40}$-$(RPRPP)_5$) exhibit a larger deviation between these two curves compared to the lowly charged peptides ($U_{40}$-$(PPRPP)_5$)

##### C. The relationship between viscosity and density

In both single- and two-component condensates from our simulations, viscosity is consistently positively correlated with the number density of monomers, i.e., $\rho \uparrow \eta \uparrow$. On a log-log plot, this relationship follows a power law: $\eta = A\rho^{\alpha}$. For GY23 condensates, which lack charged monomers in the HPS model, $\alpha \approx 3$. In contrast, RNA-peptide condensates exhibit $\alpha \approx 2$,

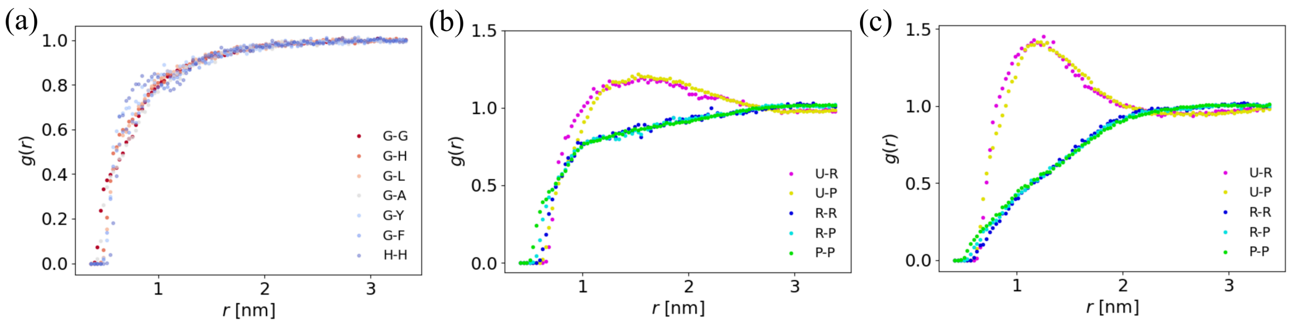


FIG. S1. Radial distribution functions $g(r)$ of residue pairs. (a) $g(r)$ for the first seven monomer pairs of WT GY23, ordered by the number of pairs within a distance of 1 nm. (b) $g(r)$ for residue pairs in $U_{40}$-(PPRPP)$_5$ (c) $g(r)$ for residue pairs in $U_{40}$-(RPRPP)$_5$.

with highly charged peptides showing a higher prefactor $A$, as illustrated by the two dashed lines in Fig.S2. Compared to single-component condensates, the strong electrostatic interactions in two-component condensates weaken the positive correlation between viscosity and density.

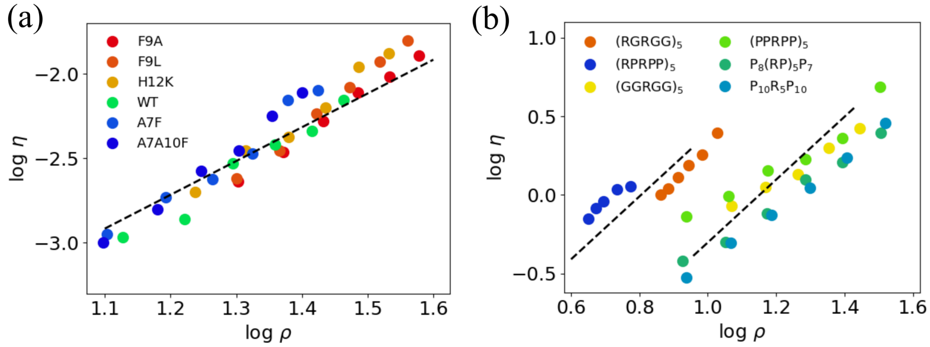


FIG. S2. The relationship between viscosity and number density in different condensates. (a) ln$\eta$ as a function of ln$\rho$ for variants of GY23; the dashed line has a slope of 3 for comparison. (b) ln$\eta$ as a function of ln$\rho$ for RNA-peptide condensates; the dashed lines have a slope of 2 for comparison.

#### D. Activation energies of different time scales of microscopic contacts

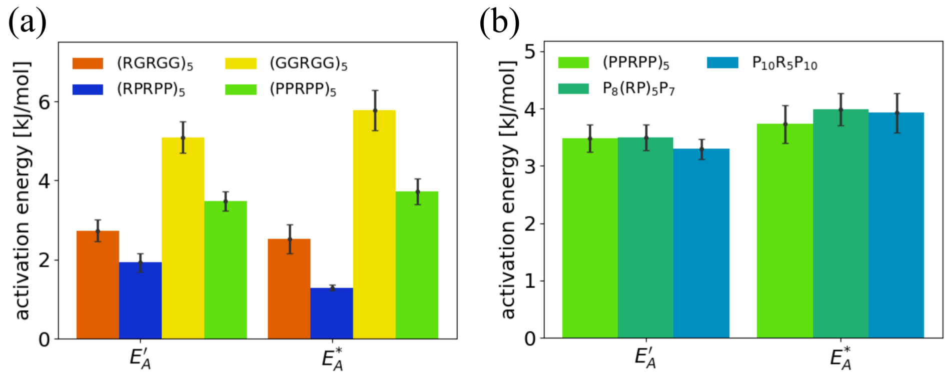


FIG. S3. Comparison of activation energy of contact lifetime $E_A'$ and $E_A^*$ for different peptide variants. (a) Variants of hydrophobicity and charge number. (b) Variants of charge pattern.

For RNA-peptide condensates, we selected the all-average contact time as a representative microscopic time scale. Since contacts between charged and neutral amino acids differ, their corresponding time scales also vary. If we shift the focus from the all-average contact group to the slower one (r-p contact for highly charged peptides and p-p for lowly charged peptides), we obtain an activation energy $E_A^*$. In most of our simulations, the value of $E_A^*$ is very close to $E_A'$ . The results for different sequence variants are shown in Fig. S3. This indicates that the activation energy derived from our 'average' contact time can effectively represent the relaxation time in the condensates."
